# Climate-driven shifts in sediment chemistry enhance methane production in northern lakes

**DOI:** 10.1101/161653

**Authors:** E.J.S Emilson, M.A. Carson, K.M. Yakimovich, J.M. Gunn, N.C.S Mykytczuk, N. Basiliko, A.J. Tanentzap

## Abstract

Freshwater ecosystems are a major source of methane (CH_4_), contributing 0.65 Pg (in CO_2_ equivalents) yr^-1^ towards global carbon (C) emissions and thereby offsetting ∼25% of the terrestrial carbon sink. Most CH_4_ emissions come from littoral sediments, where large quantities of plant material are decomposed. As climate change is predicted to shift plant community composition, and thus change the quality of inputs into detrital food webs, this can affect CH_4_ production and have far-reaching consequences for global C emissions. Here we find that variation in polyphenol availability from decomposing organic matter underlies large differences in CH_4_ production in lake sediments. Production was at least 400-times higher from sediments composed of macrophyte litter compared to terrestrial sources (coniferous and deciduous), which we link to the inhibition of methanogenesis by polyphenol leachates. Applying our estimates to projected northward advances in the distribution of *Typha latifolia*, a widespread and dominant macrophyte, we find that CH_4_ production could increase by at least 73% in the lake-rich Boreal Shield ecozone solely due to increases in this one macrophyte species. Our results now suggest that earth system models and carbon budgets should consider the effects of plant communities on sediment chemistry and ultimately CH_4_ emissions at a global scale.

**One-sentence summary:** Production of methane from lakes is at least 400-times lower when 24 sediments receive forest- as opposed to macrophyte-derived (*Typha latifolia*) litterfall.

## Introduction

Lentic freshwater ecosystems are a major source of methane (CH_4_), contributing 0.65 Pg (in CO_2_ equivalents) yr^-1^ towards global carbon (C) emissions and accounting for an estimated 6 to 16% of natural CH_4_ emissions as compared to 1% from the oceans (*1*). Freshwater CH_4_ emissions are enough to offset an estimated ~25% of the terrestrial carbon sink in CO_2_ equivalents (*2*). Within individual lakes, up to 77% of CH_4_ emissions can come from production in littoral sediments, where warm temperatures and accumulated organic matter (OM) promote methanogen activity and ebullition (*3*–*5*), and shallow waters and wave action facilitate rapid diffusion (*6*, *7*).

In northern (temperate and boreal) lakes, which account for most of the planet’s ice-free freshwater (*8*, *9*), rates of CH_4_ emission from littoral sediments are known to vary by at least three orders of magnitude (*3*), leaving considerable uncertainty to be explained in regional and global C budgets. In general, emissions are highest where littoral zones are covered with macrophytes (*3*), and plant-related CH_4_ fluxes remain one of the least-understood components of the global methane budget (*10*). Emergent aquatic plants can directly transport CH_4_ to the atmosphere through aerenchyma cells, but this cannot explain all of the variability observed within vegetated littoral zones (*7*, *11*, *12*), nor can differences in sediment temperature and OM content (*13*). Another explanation is that the activity of sediment microbial communities is inhibited, to varying degrees, by the breakdown of different OM sources (*14*), resulting in variation in the production of CH_4_ in littoral sediments.

Water-soluble polyphenol compounds from plant litter have specifically been shown to bind to and inactivate extracellular enzymes and exert toxicity in methanogens (*15*, *16*). These compounds build-up in anaerobic soils and sediments because oxygen limitation restricts phenol oxidase activity and dark conditions prevent photodegradation (*15*, *17*). In this way, the buildup of polyphenolic compounds may act similar to a ‘latch’, suppressing CH_4_ production and holding in place large quantities of C in lake sediments that would otherwise be released as CH_4_. Oxygen limitation plays a similar role in sequestering CO_2_ in peatlands by restraining phenol oxidase activity (*17*), and rates of CH_4_ production have been related to peat chemical composition (*18*).

Here we show that the production of CH_4_ in northern lakes can vary by at least 400-times because of differences in sediment chemistry related to sources of plant litterfall. We predict that sediments will differ in concentration of methanogenesis-inhibiting polyphenols according to incoming sources of OM. To test the effects of these differences in sediment chemistry on CH_4_ production in lakes, we compared natural sediments amended with OM from three widespread sources in north-temperate watersheds that would be expected to vary in polyphenol content: mixed coniferous forest litter (CON), mixed deciduous forest litter (DEC), and litter from a ubiquitous emergent macrophyte, *Typha latifolia* (TYP). The sediments were mixed at 20% OM to approximate the average concentrations found in littoral zones of northern lakes (*19*), and incubated in controlled conditions to control other effects, such as temperature, light exposure, and differences in ambient water quality, which confound observational studies. As northern watersheds are expected to experience a shift in forest composition (*20*, *21*) and an increase in emergent macrophyte growth in lakes (*22*, *23*), these findings present an additional mechanism to increased mineralization and permafrost thaw (*24*, *25*) by which climate change can enhance CH_4_ emission from northern lakes.

## Results and Discussion

After 150 days of laboratory incubation, CH_4_ production was over 400-times higher on average from *Typha latifolia* (TYP) sediments than from mixed-coniferous (CON) sediments, almost 2,800-times higher than from mixed-deciduous (DEC) sediments, and 1400-times higher than un-amended controls with 0.3% OM (CTR). In contrast, the CON and DEC treatments did not significantly differ from CTR, suggesting that methanogenesis was inhibited in the sediments amended with forest litter (Fig. 1). Our estimated CH_4_ production rates for a 150-day growing season ranged from averages of 2.63 mg m^-2^ to 7.22 × 10^3^ mg m^-2^ amongst the DEC-, CON-, and TYP-amended sediments. These production rates were comparable on a per-area basis to the range and variability of emissions measured in-situ in littoral zones of northern lakes (*3*), reflecting the close relationship between production and emission in shallow waters (*7*). We also found comparable patterns when repeating the experiment with sediments of 10 and 40% OM (Supplementary Fig. S1). A lack of differences in CO_2_ production rates amongst the amended sediments further suggested that inhibition of methanogenesis and not microbial activity in general was responsible for variation in CH_4_ production (Supplementary Fig. S2).

**Fig. 1:**
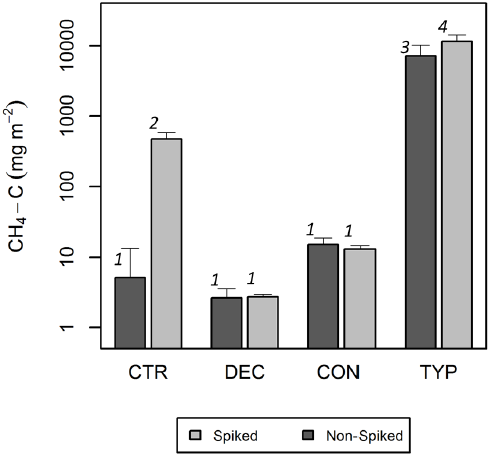
CH_4_ production in amended sediments. Production over a 150-day growing season is orders of magnitude higher in sediments amended with 20% organic matter from emergent macrophyte (*Typha latifolia*; TYP) litter than deciduous (DEC) or coniferous (CON) forest litter. CH_4_ production increases further with addition of methanogen-rich sediment (i.e. spiked-treatments) only in control (CTR) and TYP sediments. Different numbers (1-4) represent significant differences (*p* < 0.05) among amendments (ANOVA *F*_7, 24_ = 39.47). Results are shown on a log scale because of large differences between TYP and the other amendments, and error bars represent standard errors in production estimates.

We took two approaches to test the hypothesis that inhibition of methanogenesis was occurring in the lake sediments amended with forest-derived OM (CON-and DEC-treatments). Firstly, we measured the relative abundance of methanogens using qPCR targeting the *mcrA* gene and found on average 1.72× 10^2^ and 1.33 × 10^4^ fewer *mcrA* copies in the CON and DEC sediments, respectively, compared to the TYP sediments (Fig. 2). These relative abundances mirrored patterns of CH_4_ production in Fig. 1, suggesting that suppression of methanogen growth was related to decreased production of CH_4_. Although relative abundance of the *mcrA* gene that we assayed does not entirely equate with specific activity of methanogen communities, there is strong evidence linking it with CH_4_ production both here (i.e. Figs. 1-2) and in previous studies (*26*, *27*).

**Fig. 2:**
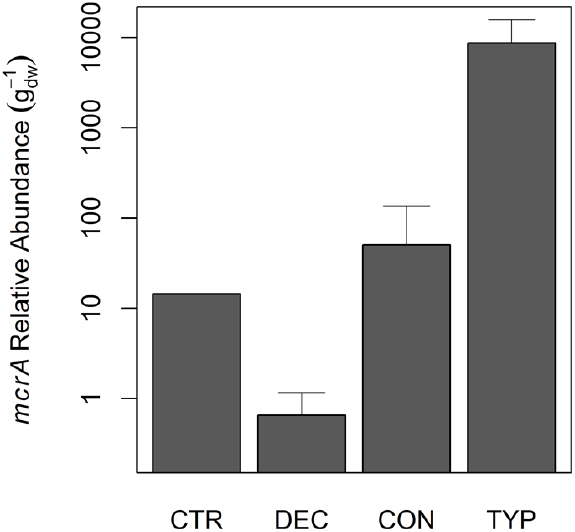
Relative abundance of *mcrA* gene copies in amended sediments. Relative abundance is orders of magnitude higher in sediments amended with emergent macrophyte (*Typha latifolia*; TYP) litter than deciduous (DEC) or coniferous (CON) forest litter and mirrors CH_4_ production in Fig. 1. DNA was pooled across replicates (n = 4 per %OM treatment) and expressed per gram dry-weight (g_dw_) of sediment normalized for extraction yield determined by qPCR. Error bars for amendments represent standard error across %OM treatments.

The second approach we took to test for inhibition of methanogenesis was to conduct a parallel set of incubations where we added a small quantity of a methane-rich sediment ‘spike’ to our treatments at the start of the experiment. Concurrent with our hypothesis of inhibition by plant-derived compounds, there was no change in CH_4_ production in the DEC or CON sediments with the spike added, but CH_4_ production doubled in the TYP sediment, and increased most strongly in the un-amended control sediments (Fig. 1). The inhibition of CH_4_ production in sediments composed of forest-derived compared to macrophyte-derived OM now offers a new mechanism to explain previously described observations in lakes wherein most of the CH_4_ emissions come from littoral zones covered with macrophytes (*3*).

Measurements of the biochemical composition of decomposing OM support our conclusion that the inhibition of methanogenesis was caused by polyphenols from the forest-derived OM. Fluorescence excitation-emission matrices of OM in sediment porewater across all the treatments revealed the presence of a protein-like fluorescence component that was associated with water-soluble polyphenolic leaf leachates (*28*, *29*), in addition to the ubiquitous tryptophan-and tyrosine-like components (Supplementary Fig. S3). Relative concentration of this water-soluble polyphenol component was lowest in the porewater of the TYP sediments, highest in the DEC sediments, and undetectable in the un-amended CTR sediments. We further found that CH_4_ production decreased with relative polyphenol concentration across all the amended sediment types and OM concentrations, suggesting that suppressed methanogenesis in CON and DEC sediments was related to water-soluble polyphenols (Fig. 3). These polyphenol leaf leachates were likely inhibiting methanogenesis by reducing enzyme and methanogen activity through direct toxicity (*16*, *30*), pH depression, and/or other chemical effects (*15*, *31*). Reduction of methanogenesis can also occur through increased availability of thermodynamically-favourable pathways in sediments (e.g. sulfate reduction), but we did not detect the presence of sulfate reducing bacteria in the sediments (below PCR detection limits; Supplementary Table S1) and so it is likely that sulfate was limiting and/or depleted during the 150-day incubation (*30*).

**Fig. 3:**
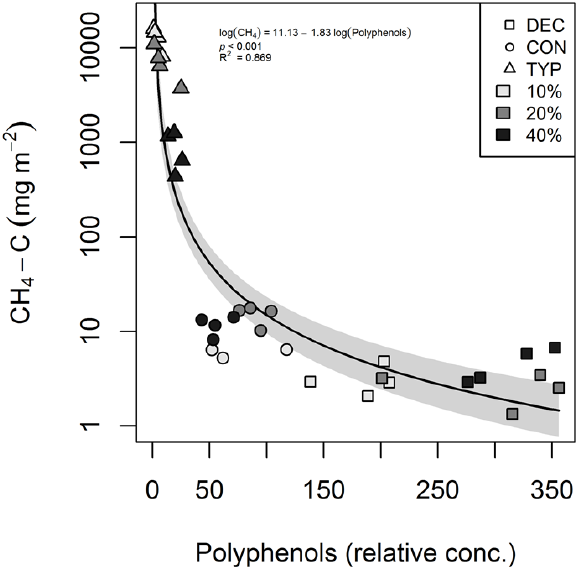
CH_4_ production in sediments declines with polyphenols. The relationship is shown across OM amendment type (DEC, CON, and TYP) and concentrations (10, 20, 40%). Concentrations of polyphenols are relative and determined from fluorescence excitation-emission spectroscopy.

As sediments amended with TYP produced so much more CH_4_ than forest litter (CON and DEC), our findings may have far-reaching implications for global carbon cycling. For example, species distribution models (SDMs) predict more favourable climatic conditions for the growth of TYP and other emergent macrophytes in boreal lakes in the coming decades (*23*, *32*). To consider the implications for CH_4_ emissions, we overlaid SDMs produced by Natural Resources Canada using published methods (*21*) onto the Boreal Shield, an ecozone with relatively homogenous underlying geology and plant communities similar to those in our incubations. By then relating projected occurrence to colonization of suitable lake habitat, we found that the number of lakes likely to be colonized by *T. latifolia* could increase by 1.7 to 2.5 times between 2041 and 2070 (Supplementary Table S2). Assuming no changes other than macrophyte spread, we estimated that the increase in *T. latifolia* alone could elevate CH_4_ production across Boreal Shield lakes by at least 73% during a 150-day growing season (Supplementary Table S2, Fig. 4). Of course, these estimates are heavily caveated by several assumptions. For example, climate-driven changes in other factors, such as temperature, oxidation potential, and increased forest litterfall production, will certainly influence CH_4_ production from lake sediments, and all production may not necessarily result in emissions (*1*). We have also not accounted for aerenchymal transfer, which may further enhance emissions where TYP is present, nor differential mixing of forest-derived OM in sediments resulting from expected shifts in deciduous forest cover (*33*). However our rough calculation is intended to emphasise that lake sediment chemistry is sufficiently important that it should be considered in earth system models, or at the very least in lake carbon budgets (*34*).

**Fig. 4:**
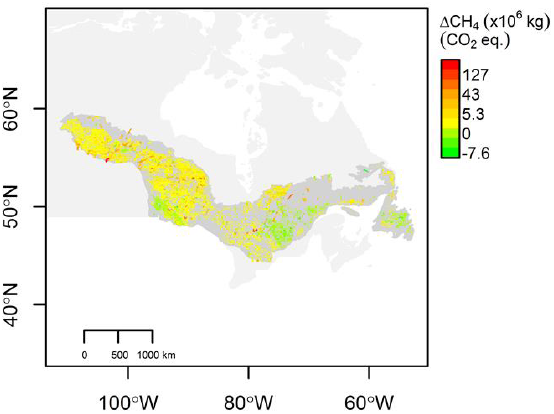
Predicted increase in CH_4_ production across the Boreal Shield. An increase of at least 73% is predicted because of the greater probability of occurrence of *Typha latifolia* alone. Estimated change in production is shown here as change in total kg of CH_4_ (CO_2_ equivalents) from current (1971-2001) to future (2041-2070) over a 150-day growing season in each lake under Composite-AR5 RCP 4.5 climate scenario.

Methane production in freshwater ecosystems has recently been recognized as an important component of global C cycles (*2*). Here we have discovered a new mechanism by which plant-related shifts in sediment chemistry under a changing climate can increase methane production in lakes. This mechanism can account for the observed variability in CH_4_ emission that has been reported both across and within lakes (*2*, *3*), and should enable more precise models and C budgets in northern watersheds.

## Materials and Methods

### Experimental Design

We amended natural sediments with three different sources: senescent coniferous (CON) and deciduous (DEC) litterfall from a transitional/mixed forest stand (Central Ontario: 44°7'22.3"N,79°30'23.7"W), and senescent *Typha latifolia* (TYP) from Ramsey Lake (in Sudbury, Canada: 467°28'19.8"N 80°58'19.2"W). The CON mix consisted of *Pinus resinosa* and *Pinus strobus*, and the DEC mix consisted primarily of *Acer rubrum*, *Acer saccharum*, *Betula* spp., *Populus tremuloides*, *Ulmus americanum*, *Quercus rubra*, and *Quercus alba*. All OM was oven dried for 12 h at 60 °C, ground, and sieved to retain only the fine particulate organic matter (FPOM) fraction (≤ 1mm).

We mixed the FPOM with a “base inorganic sediment” (0.3% OM, determined by loss-on-ignition at 500°C for two hours) to create final OM concentrations (by dry-weight) of 20% across the three amendments (CON, DEC, TYP). We used 20% to approximate typical OM concentrations found in littoral zones of northern lakes (*19*) (and confirmed in a nearby lake (*35*)) but we also measured CH_4_ production with 10 and 40% OM to confirm similarity of patterns across conditions. The base sediment was collected from the shoreline of Geneva Lake (near Sudbury, Canada: 46°45'27.2"N, 81°33'19.8"W) away from *T. latifolia* beds and direct inputs of forest-derived OM and was sieved to exclude particles larger than 2 mm. We distributed the mixed sediments into 250 mL mason jars equipped with rubber septa, with four replicate jars per each %OM and amendment type combination. An estimated 70% of methane production occurs in the top 5 cm of saturated soils (*36*), so we filled the jars to a depth of 4.5 cm (allowing room for expansion), before saturating them with TOC-scrubbed A10 MilliQ water (EMD Millipore Corp., Darmstadt, Germany). We also created replicated control jars containing only base sediment, otherwise constructed and treated in the same manner.

We duplicated the 20% OM experimental setup with a “methane-rich spike”. The spike consisted of replacing 5% of the base sediment with sediment from the top 5 cm of a littoral site in Ramsey Lake previously known to have high rates of methane production. Amendments of CON, DEC, TYP were adjusted for the 2.8% OM content of the spike sediment to ensure final OM concentrations of 20% (dry-weight).

### CH_4_ and CO_2_ Production

We incubated the sediments and periodically collected headspace samples to measure CH_4_ and CO_2_ production over 150 days, representative of the length of a growing season in the Boreal Shield. The sediments were incubated in a BioChambers SPC-56 growth chamber in the dark at 20.5°C. At the start of the incubations, headspace air in each jar was replaced four times with N_2_ using a vacuum manifold to ensure anaerobic conditions and removal of atmospheric CO_2_ and CH_4_. We collected headspace gas on days 5, 10, 15, 31, 60, 91, 120, and 150 by homogenizing 10 mL of N_2_ into headspace prior to extracting a 10 mL gas sample by syringe. The total volume removed was quantified and used to correct headspace volume throughout the incubation. Both CH_4_ and CO_2_ were detected as CH_4_ using a SRI 8610C gas chromatograph (0.5 mL sample loop, 105°C column temperature), and production was calculated at the end of 150 days, adding back the portions that were removed and expressing totals as mg m_-2_ of dry sediment given an area of 28.3 cm^2^ for each jar.

### Relative Polyphenol Concentration

To measure relative polyphenol concentration, we collected porewater from each jar after the 150-day incubations and filtered the samples through 0.5 μm glass fiber filters. Samples were acidified to pH < 2 with HCl, and stored in airtight vials at ∼4°C. Fluorescence EEMs (excitation-emission matrices) were generated using an Agilent Cary Eclipse Fluorescence Spectrophotometer in ratio (S/R) mode with a 1 cm path-length cuvette. EEMs were generated from excitation and emission intensities (EX: 250 to 450 nm in 5 nm steps, EM: 300 to 600 nm in 2 nm steps) that were adjusted for inner-filter effects with absorbance as measured with an Agilent Cary 60 UV-Vis Spectrophotometer. All EEM sample correction and PARAFAC modelling was done in Matlab R2015b according to the methods outlined in ref. (*37*).

Five PARAFAC components were validated by a split-half method (*37*), explaining 98.7% of the variation in the EEMs. Components C1 and C2 were comparable to common humic-like components, with maximum excitation/emission intensities of (310/414 nm) and (345/462 nm) respectively. C3 was similar to the common tryptophan protein-like component (280/354 nm), and C4 the common tyrosine protein-like component (270/306 nm). Component C5 (275/318 nm) was identified as a protein-like component that is associated with leaf litter polyphenol leachates (*28*, *29*) (Supplementary Fig. S3). Therefore, relative polyphenol leachate concentration was estimated as the product of proportional C5 fluorescence and total dissolved organic carbon (DOC) concentration in sediment porewater, as measured on a Shimadzu TOC-5000A in FPOC mode.

### Methanogen Suppression

To compare the relative abundance of methanogens between samples, DNA was first extracted in duplicate using the MoBio PowerSoil kit (MoBio, Carlsbad, CA, USA). qPCR was then carried out in triplicate on pooled DNA extractions to characterize better the communities from the 4 sediment replicates of the CTR and each OM type mixed at 10, 20 and 40% concentration. The *mcrA* gene was targeted using mlasF (5’-GGYGGTGTMGGDTTCACMCARTA-3’) and mcrA-rev (5’ CGTTCATBGCGTAGTTVGGRTAGT-3’) primers as in ref. (*26*). Reaction conditions were: a 5-minute initial denaturation at 95 °C, followed by 45 cycles of 95 °C for 15 seconds, 55 °C for 30 seconds, and 72 °C for 30 seconds. Then a final denaturation for 1 minute at 95 °C, 30 seconds at 42 °C and 95 °C for 30 seconds. The qPCR was done using Biorad’s iTaq Universal SYBR Green Supermix on an Agilent Technologies Stratagene Mx3005P. A standard curve was generated by serially diluting an extracted band from amplified eDNA and run in triplicate along with the samples generating an R^2^ = 0.999 and efficiency of 97.2%. Dissociation curves indicated a pure product, which was confirmed on a 1.5% agarose gel. eDNA was quantified and purity was checked (260/280 nm ratio) using a Take3 spectrophotometry system on a Synergy HI microplate reader (BioTek, Winooski VT, USA). Dissociation curves indicated a pure product, which was confirmed on a 1.5% agarose gel. The results were calculated by averaging the triplicate Ct values, and abundances were standardized relative to the control and expressed per dry weight of sediment normalized for extraction efficiency. Suppression of methanogens could also be caused by sulfate reducing bacteria (SRB). To test for this, we used PCR to target the SRB-specific *dsr*A gene and 16S rRNA deep sequencing to evaluate their abundance (see Supplementary Methods S1).

### Ecosystem-scale Emissions

We estimated the impact of increased *T. latifolia* occurrence on CH_4_ production during the growing season by applying our estimated production rates (mg m_-2_) to current and projected aerial cover (m^2^) for Boreal Shield lakes. Surface areas were obtained from the Global Lakes and Wetlands Database (GLWD) (*8*) for waterbodies between 0.1 and 1,000 km^2^ in size and located within the Boreal Shield (spatially delineated by the National Ecological Framework for Canada (*38*) as an area of 1.8 million km^2^ located between ca. 45°N and 60°N characterized by underlying Precambrian bedrock). For each lake, we extracted current and projected probability of occurrence of *T. latifolia* from Natural Resources Canada MaxEnt raster data, which was developed by combining species occurrence data with actual climate data (for current estimates) and with climate models (GCMs) (*21*, *32*). We used projected MaxEnt occurrences for the timeframe of 2041-2070 incorporating uncertainty by using five climate models [canESM2, hadGEM2-ES, CESM1(CAM5), MIROC-ESM-CHEM, and composite-AR5] each with three future emission scenarios (RCP 2.6, 4.5 and 8.5).

We then used the range in the probability of occurrence data to estimate a range in projected suitable habitat and thus proportional coverage within Boreal Shield lakes. Suitable emergent macrophyte habitat (shallow littoral) areas are not widely available, so we used published regressions indicating a maximum of 28% of lake area to be covered by emergent macrophytes, on average, for Boreal lakes in our size range (*39*). We then estimated coverage as the product of the probability of occurrence and 28% of the total lake areas that were widely available across the Boreal from the GLWD. Thus, a probability of occurrence of 1 meant all suitable habitat, or 28% of total lake area, was covered in a given lake. We then calculated total CH_4_ production as the product of the rate of production (mg m^-2^) in our incubation study and coverage by *T. latifolia* (m^2^, current and projected), propagating uncertainty from climate models and scenarios along with variation in our CH_4_ production estimates. Estimates were scaled up to 100% of the sediment profile assuming our 5 cm surficial sediments represented 70% of total production (*36*) and presented in CO_2_ equivalents (1 kg CH_4_ = 25 kg CO_2_) to maintain consistency with global emission estimates in ref. (*2*).

### Statistical Analysis

To compare production rates across OM type, we performed a one-way ANOVA in R 3.3.0 (*40*). The ANOVA included the effect of type and its interaction with the methanogen spike with the baseline (intercept) group adjusted to compare significance among groups. The ANOVA was repeated for 10, 20, and 40% OM separately. We then fit a log-log model in R 3.3.0 (32) to test for an effect of relative polyphenol concentrations on CH_4_ production. All spatial analyses were also done with R v. 3.3.0 (*40*).

## Supplementary Materials

Methods S1. PCR detection and 16S rRNA sequencing libraries for sulfate reducing bacteria.

Fig. S1. CH_4_ production in 10 (A) and 40% (B) amendments.

Fig. S2. CO_2_ production in amended sediments.

Fig. S3. Five dissolved organic matter PARAFAC components identified in sediment porewater.

Table S1. PCR results for sulphate reducing bacteria (SRB).

Table S2. Production estimates from Boreal Shield lakes for a 150-day growing season.

## Acknowledgements

We acknowledge the support of staff and colleagues at the Living with Lakes Centre, Laurentian University, in Sudbury Canada. We also thank John Pedlar and Dan McKenney at Natural Resources Canada for providing spatial MaxENT data.

## Funding

Support was provided by NERC Standard Grant NE/L006561/1.

## Author contributions

ESE, MC, JG, NM, NB, and AJT conceived the study. ESE, MC, and KY collected the data, and ESE analyzed the data and wrote the manuscript with input from all authors.

## Competing interests

The authors declare that they have no competing interests.

## Data and materials availability

All data are present in the paper and/or the Supplementary Materials. Additional data related to this paper may be requested from the authors. All SRB related sequences have been deposited in the NCBI Sequence Read Archive under BioProject PRJNA347436.

